# AlphaFold Models of Small Proteins Rival the Accuracy of Solution NMR Structures

**DOI:** 10.1101/2022.03.09.483701

**Authors:** Roberto Tejero, Yuanpeng J. Huang, Theresa A. Ramelot, Gaetano T. Montelione

## Abstract

Recent advances in molecular modeling using deep learning have the potential to revolutionize the field of structural biology. In particular, AlphaFold has been observed to provide models of protein structures with accuracy rivaling medium-resolution X-ray crystal structures, and with excellent atomic coordinate matches to experimental protein NMR and cryo-electron microscopy structures. Here we assess the hypothesis that AlphaFold models of small, relatively rigid proteins have accuracies (based on comparison against experimental data) similar to experimental solution NMR structures. We selected six representative small proteins with structures determined by both NMR and X-ray crystallography, and modeled each of them using AlphaFold. Using several structure validation tools integrated under the Protein Structure Validation Software suite (PSVS), we then assessed how well these models fit to experimental NMR data, including NOESY peak lists (RPF-DP scores), comparisons between predicted rigidity and chemical shift data (ANSURR scores), and ^15^N-^1^H residual dipolar coupling data (RDC Q factors) analyzed by software tools integrated in the PSVS suite. Remarkably, the fits to NMR data for the protein structure models predicted with AlphaFold are generally similar, or better, than for the corresponding experimental NMR or X-ray crystal structures. Similar conclusions were reached in comparing AlphaFold2 predictions and NMR structures for three targets from the Critical Assessment of Protein Structure Prediction (CASP). These results contradict the widely held misperception that AlphaFold cannot accurately model solution NMR structures. They also document the value of PSVS for model *vs*. data assessment of protein NMR structures, and the potential for using AlphaFold models for guiding analysis of experimental NMR data and more generally in structural biology.

## Introduction

Recent advances in protein structure prediction based on deep learning from experimental protein structure data have the potential to revolutionize structural biology. Building on advances in attention-based machine learning (Vaswani et al., 2017; Huang et al., 2019), contact prediction based on sequence covariance (Marks et al., 2011; Morcos et al., 2011; Marks et al., 2012; Ovchinnikov et al., 2015; Ovchinnikov et al., 2016; Buchan and Jones, 2018), the massive and still growing databases of genomic sequence data, and the rapidly growing Protein Data Bank of experimental protein structures, these methods are being recognized as major advances enabling new structural biology research (Jones and Thornton, 2022).

In the 2020 Critical Assessment of Protein Structure Prediction (CASP14), the DeepMind AlphaFold2 (AF2) deep learning method (Jumper et al., 2021a; Jumper et al., 2021b) demonstrated outstanding performance in blind predictions of protein structure, delivering excellent structural matches to experimental models derived from X-ray crystallography, NMR and cryoEM data, over a wide range of target difficulty (Kryshtafovych et al., 2021). These AlphaFold2 model predictions had an unprecedented high accuracy, assessed by backbone atomic coordinate global distance test (GDT_TS) scores. On 96 CASP14 targets AF2 models had a mean GTD_TS of 0.88 ± 0.1, corresponding to a backbone atom root-mean-squared deviation (RMSD) between predicted and experimental protein structures of about 1.5 Å (Kryshtafovych et al., 2021). Buried sidechain conformations in these blind predictions of protein structure are also generally a remarkably good match between the predicted model and experimental structure (Pereira et al., 2021). Soon afterward, the related RosettaFold (Baek et al., 2021b) method also demonstrated excellent modeling accuracy on natural proteins, and was found to be particularly successful in modeling *de novo* designed proteins. These results have opened the door to innovative *de novo* protein design approaches using these platforms (Anishchenko et al., 2021). In addition, it was also quickly recognized that the structures of protein-protein complexes and multimeric assemblies can often be reliably modeled using modified approaches with these same AI platforms (Baek et al., 2021a; Baek et al., 2021b; Evans et al., 2021; Humphreys et al., 2021; Colman et al., 2022; Mondal et al., 2022). While challenges remain, particularly for dynamic protein systems and complex multiprotein assemblies, these methods are already having a major impact on structural biology.

These advances are particularly relevant for structural studies of proteins using NMR data. In the 2017 CASP13 blind protein structure prediction experiment, we organized a “NMR-guided prediction” challenge for the CASP protein structure prediction community called CASP-NMR (Sala et al., 2019). In this project we provided 13 simulated and real NMR data sets for 10 small (80 to 326 residues) proteins, including interatomic contacts obtained (or, for simulated data, obtainable) from NOESY experiments and backbone dihedral restraints obtained (or obtainable) from backbone chemical shift data. These included NOESY data typical of that obtained for ^15^N,^13^C-enriched, perdeuterated proteins up to about 40 kDa, which were simulated and used to generate tables of ambiguous contacts using simple NOESY peak assignments protocols. These Ambiguous Contact Lists were provided, together with backbone dihedral angle restraints obtainable from chemical shift data, to the CASP prediction community for data-assisted prediction. Real NMR data collected for a *de novo* designed protein were also used to generate ambiguous contact tables and chemical-shift based backbone dihedral angle restraints, that were then provided to CASP13 predictor groups, including one set of (ambiguous) experimental NMR-based contacts in which only backbone resonance (no sidechain) assignments were available. The CASP community was then challenged to use these data to “guide” blind protein structure predictions. These predictions were compared to NMR-based models generated from these data by experts using conventional methods, or against the NMR data. Remarkably, several CASP13 prediction groups provided models that matched the reference structures and/or fit these NMR data even better than the models generated by conventional expert NMR structure analysis. Notable among these top-performing NMR-guided prediction groups were methods using NMR-guided MELD (Robertson et al., 2019) and NMR-guided Rosetta (Kuenze and Meiler, 2019) methods. Amazingly, some other CASP13 prediction groups provided pure prediction models, which did not use the NMR data at all, that also matched the reference structures better than the models generated with the data by expert data analysis (Sala et al., 2019). Among the top performing groups in this category were machine learning methods including AlphaFold2.

Even more exciting results came in late 2020 from the CASP14 blind protein structure prediction experiment. In CASP14, the next-generation AlphaFold2 methods demonstrated outstanding performance in protein structure prediction (Kryshtafovych et al., 2021; Pereira et al., 2021). Interesting results were observed for three CASP14 targets for which NMR data were available (Huang et al., 2021). For two of these, the pure prediction AlphaFold2 models were observed to fit real experimental NMR data as well or better than the reference structures provided by the experimental NMR groups. In a third case, target T1027, a protein exhibiting spectral properties indicating extensive conformational dynamics, the AlphaFold2 model did not fit the NMR data as well as the reference experimental NMR structure. The AlphaFold2 prediction model of a fourth NMR target, the 238-residue integral membrane protein MipA, also was an excellent fit to the experimental NMR data. This study also demonstrated how an AlphaFold2 model of CASP14 target T1029 could be used to guide reanalysis of the experimental NMR NOESY data to provide a revised experimental structure which better fits other NMR structure quality assessment metrics, including residual dipolar coupling (RDC) Q factors and ANSURR scores. The conclusions of this CASP14 - NMR study, using blind predictions for targets not made available to the prediction groups, are supported by two other recent studies of the modeling accuracy of AlphaFold, using as reference either previously deposited NMR structure coordinates (Zweckstetter, 2021), or X-ray crystal structures and experimental RDC data (Robertson et al., 2021).

The Protein Structure Validation Software suite (PSVS) (Bhattacharya et al., 2007) integrates multiple tools for protein structure validation, with a particular focus on protein NMR structure validation. PSVS provides several *knowledge-based protein structure validation* tools, including Molprobity clash and Ramachandran backbone analyses (Lovell et al., 2003; Chen et al., 2010), as well as Verify3D (Luthy et al., 1992) and ProsaII (Sippl, 1993) protein fold analysis tools. PSVS also provides a *model vs data protein structure validation* analysis using the PDBStat (Tejero et al., 2013) software for distance and dihedral angle restraint violation analysis, supporting several common distance restraint formats and providing conversion of distance and dihedral angle restraints between these formats to allow interoperability between various NMR structure modeling software packages. Recently, PSVS has been updated (version 2.0) to include additional model vs data structure validation tools, including the RPF-DP score (Huang et al., 2005; Huang et al., 2012) comparing structure models against NOESY peak list data, RDC Q factors (Cornilescu et al., 1998; Clore and Garrett, 1999) comparing models against RDC data, and the ANSURR (Accuracy of NMR Structures Using RCI and Rigidity) score (Fowler et al., 2020) comparing measures of conformational rigidity for various regions of protein structure models with metrics of rigidity based on backbone chemical shift data.

Here, we test the hypothesis, based on our experiences in CASP13 and CASP14, that small protein structures modeled using the recently released AlphaFold platform fit experimental NMR NOE, RDC, and structural rigidity (chemical shift) data as well as models generated by experts using conventional NMR data analysis methods. Six proteins solved by expert NMR spectroscopists in the course of the PSI Structural Genomics Initiative were modeled by AlphaFold and then assessed against both experimental NMR data and knowledge-based statistical metrics using the PSVS software suite. AlphaFold modeling was done by excluding information from the deposited structure itself, or from any homologous templates, as input information. *In all cases, the various PSVS structure quality scores for AlphaFold models document that these predicted structures fit the NMR data as well, or often better, than experimental structures deposited in the Protein Data Bank by expert spectroscopists*. Overall, this study demonstrates the outstanding value of AlphaFold for modeling small, relatively rigid protein structures, and for providing atomic coordinates useful in guiding analysis of experimental data.

## Methods

### NMR and X-ray structure coordinate sets and NMR data

Experimentally-determined protein structure coordinates and NMR data were taken for proteins deposited in the Protein Data Bank (PDB) by the Northeast Structural Genomics Consortium (Montelione et al., 2013). For this study we used small proteins solved by both NMR and X-ray crystallography methods, and for which both nearly complete resonance assignment and NOESY peak list data are publicly available from the *NESG NMR / X-ray Pairs* web site (Everett et al., 2016) (https://montelionelab.chem.rpi.edu/databases/nmrdata/). For three of these proteins, ^15^N-^1^H residual dipolar coupling data are also available from this site. The proteins ranged in size from 58 to 158 residues (excluding short hexa-His purification tags). These same atomic coordinates and most of these NMR data are also available in the PDB and/or the BioMagResDataBase (BMRB). In addition, structures of these same six proteins that have been energy-refined using the NMR-restrained Rosetta refinement protocol (Mao et al., 2014) were also obtained from the *NESG / NMR X-ray Pair* web site. For the two structures for which RDCs were used in the original structures deposited in the PDB (SrG115C and RpR324), the Rosetta refinement was carried out with these RDC data. For a third target (SgR209C) RDC data is also available in the NESG database, but it was obtained only after the structure was deposited in the PDB, and was not used in the original structure determination nor in the Rosetta refinement.

### Solution NMR structure determinations

For two cases required for this study, target protein structures were redetermined from the original NMR data following the standard methods of the NESG consortium (Liu et al., 2005; Montelione and Szyperski, 2010). The solution NMR structure ensemble for RpR324 was recalculated using previously described NMR data (Ramelot et al., 2012), including the resonance assignments, NOE, dihedral, and hydrogen bond restraints of PDB entry 2LPK, but excluding the RDC data, using CYANA (Güntert and Buchner, 2015) followed by refinement with CNS in explicit water. The structure of target SgR209C, PDB entry 2LO6 determined without RDC data, was redetermined using the NOE, dihedral, and hydrogen bond restraints from PDB entry but also including ^15^N-^1^H RDC RDC data measured on samples partially aligned in polyacrylamide stretched gel (PAG) and polyethylene glycol (PEG) alignment media that were not used in the original structure determination. These RDC data were also obtained from the *NESG NMR /X-ray Pairs* web site. RDC data were manually assessed and excluded if they had peak overlap in the ^1^H-^15^N 2D plane or were in disordered regions of the protein. This resulted in 65/90 PAG and 69/103 PEG RDCs used in the refinement, carried out using the same standard methods outlined above for RpR324.

### AlphaFold (AF) modeling

AF modeling was carried out using the AF-multimer software (Evans et al., 2021) installed on the NPL cluster in the Center for Computational Innovation at Rensselaer Polytechnic Institute. The system has 40 nodes each with 2x 20 core 2.5 GHz Intel Xeon Gold 6248 CPUs and 8x NVIDIA Tesla V100 GPUs, with 32 GiB HBM, with 768 GiB RAM per node. This version of AF was trained using the PDB database of April, 2018 and did not use any NMR structures in the training data (Jumper et al., 2021a). PDB structures deposited after November 2005, including the X-ray crystal and NMR structures of the query target proteins structures themselves and the structures of any homologs detected with HMMPred (Soding et al., 2005) were excluded as modeling templates. Hydrogen atoms were added by the AF prediction pipeline to each model using restrained refinement of the H atom positions with Amber (Case et al., 2021) with fixed heavy atom coordinates. The AlphaFold model was represented by five top-scored conformations along with estimates of prediction reliability (pLDDT), as described elsewhere (Jumper et al., 2021a).

### Knowledge-based protein structure model validation

All structure quality statistical analyses were performed using the Protein Structure Validation Software (Bhattacharya et al., 2007) (PSVS ver 2.0-pre) (https://montelionelab.chem.rpi.edu/PSVS/PSVS2/). PSVS runs a suite of knowledge-based software tools including PDBStat (ver 5.21.6) (Tejero et al., 2013), ProCheck (ver 3.5.4) (Laskowski et al., 1993), MolProbity (mage ver 6.35.040409) (Chen et al., 2010), and an implementation of the algorithms of FindCore2 (Snyder et al., 2014) coded in PDBStat. The structure validation scores of these programs were used to calculate a normalized Z score relative to mean values and standard deviations obtained for a set of 252 reference X-ray crystal structures of < 500 residues, resolution < 1.80 Å, R-factor < 0.25 and R-free < 0.28; positive Z scores indicate ‘better’ scores.

### RDC Q score analysis

^15^N-^1^H residual dipolar couplings D_calc_ were calculated from model structures by single-value decomposition of the Saupe matrix (Losonczi et al., 1999) using PDBStat (Tejero et al., 2013), called from PSVS ver 2.0. Residual dipolar coupling Q factors were analyzed by PDBstat using both of the following methods. The most commonly used RDC-fit score Q1, described by Cornilescu et al (Cornilescu et al., 1998) is

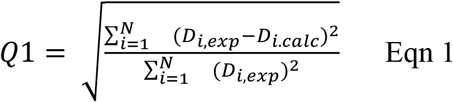

where D_exp_ and D_calc_ are the measured and calculated values of the RDC, and N is the number of RDCs assessed. In addition, we also assessed models using RDC-fit score Q2, described by Clore et al (Clore and Garrett, 1999) and used by the DC: Servers for Dipolar Coupling Calculations (https://spin.niddk.nih.gov/bax/nmrserver/dc/svd.html).

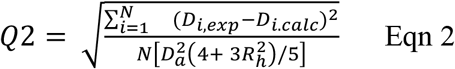

where *D_a_* is the axial component, and *R_h_* is the rhombic component, of the orientation tensor. The Q2 factor is preferable in case of a limited RDC sampling over all possible orientations (Clore and Garrett, 1999).

### RPF-DP scores

RPF-DP scores are a set of fast and sensitive structure quality assessment measures used to evaluate how well a 3D structure model fits with NOESY peak and resonance assignment lists, and hence to assess the accuracy of the structure (Huang et al., 2005; Huang et al., 2012). RPF-DP scores provide a type of NMR “R-factor”, in which models are compared against unassigned NMR NOESY peak list data. RPF-DP scores were computed with the program RPF, called from PSVS. An RPF server is also available online at https://montelionelab.chem.rpi.edu/rpf/. RPF-DP metrics have been described previously (Huang et al., 2005; Huang et al., 2012; Huang et al., 2021), but as they play a key role in this work, we provide a brief overview of these model vs data structure quality assessment metrics here. Additional details are provided in the original paper (Huang et al., 2005).

The RPF-DP score algorithm is outlined schematically in Figure 1, adopted from Huang et al (Huang et al., 2005; Huang et al., 2012; Huang et al., 2021). Nodes represent all protons listed in the resonance assignment table. Edges connect the nodes and represent all potential associated NOEs from the NOESY peak lists, within a chemical shift match tolerance. In constructing the ambiguous NOE network G_ANOE_ (shown on right side of Figure 1), each NOESY cross peak (p) may be ambiguously assigned to one or more proton pairs, as determined by chemical shift degeneracies and match tolerances. The true NOE network, G_NOE_, corresponding to the true 3D structure(s), is a subgraph of G_ANOE_. Given complete NOESY peak lists and resonance assignments, for each NOESY cross peak p, at least one of its possible proton pair assignments has a corresponding edge in G_NOE_. For each structure model (shown on left side of Figure 1), a distance network G is calculated, from the summation distances (sum of inverse sixth powers of individual degenerate proton-proton distances), assuming uniform effects of nuclear relaxation processes. Nodes of G are connected by an edge if the corresponding interproton summation distance in the model structure is ≤ d_NOE_max_, where d_NOE_max_ is the (estimated) maximum distance detected in the NOESY spectrum. Summation distances are used to address the lack of stereospecific assignments of prochiral methylene proton pairs, sets of protons that are degenerate (e.g. the three hydrogens of a methyl group, degenerate methylene protons, or degenerate resonances of Tyr or Phe), or combinations of these kinds of ambiguities (e.g. for prochiral isopropyl methyl groups of Leu or Val for which stereospecific assignments are not available). The default upper-bound observed distance, d_NOE_max_, used in these metrics is 5 Å, but this can also be calibrated from the NOESY data. For models derived by X-ray crystallography, protons are added with ideal covalent geometry using the program Reduce v2.14 (Chen et al., 2010).

**Fig. 1.**
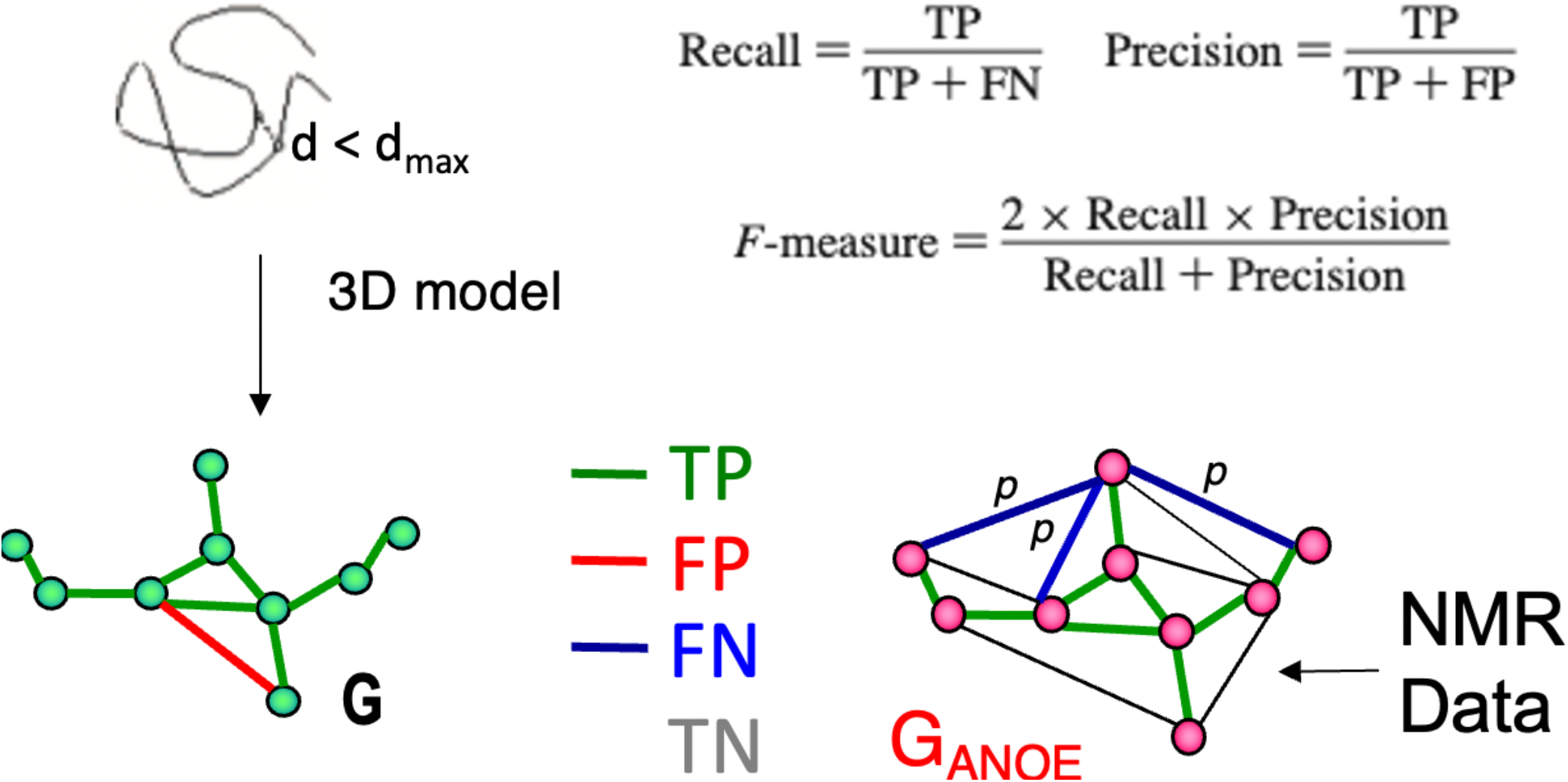
Schematic description of RPF-DP scores. In this analysis, the graph G with nodes corresponding to all ^1^H’s and edges representing all short (e.g., < 5 Å) ^1^H-^1^H distances in a structure model (left), is compared with a graph G_ANOE_ (right), in which nodes again correspond to all ^1^H’s and edges describe all possible assignments for each NOESY cross peak. True positives (TPs) are edges common to both G and G_ANOE_, false positives (FPs) are edges present in G but not in G_ANOE_, and false negatives (FNs) are the set of edges in G_ANOE_ representing the multiple possible assignments of a NOESY cross peak, none of which are present in G. These metrics are used to compute recall (R), precision (P), and F-measure as shown in the figure and outlined in the Methods Section. The F-measure is the harmonic mean of the recall and precision. The Discriminating Power (DP) is a normalized F-measure corrected to account for the F-measure expected for a random-coil chain (DP = 0) and the best F-measure possible considering the completeness of the NMR data (DP = 1.0). Since NOESY data is restricted to short distances (e.g., < 5 Å), true negatives (TNs, peaks not expected from the model and not observed in the NOESY data) can dominate these statistics and are not included in these recall, precision, and F-measure metrics. Figure and legend are adopted from (Huang et al., 2021).

As illustrated in Figure 1, proton pair short distances present in the atomic coordinates of a model structure, represented by the network G, may or may not be represented in the graphical representation of the NOESY peak list data G_ANOE_. NOESY cross peaks represented in G_ANOE_ that are consistent with the short interproton distances in the network derived from the model, G, are defined as true positives (TPs), while NOESY peaks expected from the model (edges in G) that are not observed in the data, G_ANOE_, are false positives (FPs). Since G_ANOE_ is an ambiguous network, a FN score is assigned to a NOESY peak only if none of the several possible short proton-proton distances consistent with these several possible NOESY peak assignments are observed in the structure model, represented by G. In this context, recall (R) measures the fraction of NOE cross peaks that are consistent with the query structure models, while precision (P) measures the fraction of short proton pair distances in the structure model that are observed in the NOESY peak list (i.e., in G_ANOE_), weighted by interproton distance to minimize the impact of weak NOEs arising from interproton distances near the edge of the defined distance cutoff (Huang et al., 2005). Hence “recall violations’’ are NOESY peaks (or “noise peaks”) that cannot be explained by the model and resonance assignments, and “precision violations” are short distances in the model that are not supported by NOESY data, which result from overpacking, incorrect sequence-specific assignments, or from exchange broadening. The F-measure is the harmonic mean of the recall and precision. Equations used to calculate recall (R), precision (P), and F-measure (F, also called the performance) are shown in Figure 1. The DP score is a scaled F-measure that accounts for lower-bound and upper-bound values of the F-measure. The lower-bound of F(G) is estimated by F(G_free_), where G_free_ is a distance network graph computed from interproton distances in a freely rotating polypeptide chain model, as described by Flory and co-workers (Flory, 1969), and the upper-bound of F(G) is determined by assessing the completeness of the NOESY peak list data for the 3- and 4-bond connected H atoms which all have interproton distances < 5 Å.

RPF-DP scores can be calculated either for individual models from an ensemble of conformations, and averaged, or using average interproton distances across the ensemble. The ensemble DP score (<DP>) is usually 10 - 15% higher than average of individual DP scores (DP_avg_). However, when the conformational ensemble is more diverse, larger differences between DP_avg_ and <DP> are observed. In various studies (Huang et al., 2005; Raman et al., 2010; Huang et al., 2012; Rosato et al., 2012; Rosato et al., 2013; Rosato et al., 2015; Sala et al., 2019; Huang et al., 2021), structures within 2.0 Å RMSD of the corresponding expert-derived “correct” structure have been observed to have <DP> scores > 0.70 for NMR ensembles, and DP_avg_ scores > 0.60 averaged over the individual conformers.

### ANSURR scores

The Accuracy of NMR Structures Using RCI and Rigidity (ANSURR) method provides an independent assessment of model accuracy by comparing protein flexibility computed from backbone chemical shifts with protein flexibility predicted with a graph theory based measure of structural rigidity (Fowler et al., 2020). ANSURR provides two measures of similarity between these measures, a correlation score (corr) which assesses the correlation between peaks and troughs of observed and predicted structural flexibility along the sequence, and root-mean-squared deviation (RMSD) between these metrics. Both the corr and RMSD score are reported as a percentile score (ranging from 0 to 100). These scores were calculated using ANSURR program version 1.2.0.

### Well-defined residue ranges and Global Distance Test scores

For NMR structure ensembles, the ranges of residues that are “well-defined” were determined by standard conventions instantiated in the programs Cyrange (Kirchner and Güntert, 2011) and FindCore2 (Snyder et al., 2014). Following the recommendations of the wwPDB NMR Structure Validation Task Force (Montelione et al., 2013), residues were used in superimpositions and structure quality assessment if they are “well-defined” (i.e. well-converged) across the NMR structure ensemble and also “reliably predicted” by AlphaFold, as outline in our previous studies comparing NMR-derived and AlphaFold models (Huang et al., 2021).

GDT_TS scores were computed by the method of Zemla (Zemla, 2003; Zhang and Skolnick, 2004) using the representative “medoid” conformer (Montelione et al., 2013; Tejero et al., 2013) from each of the NMR-derived or AlphaFold conformational ensembles, superimposed using backbone C^α^ atoms within the “well defined” residue ranges:

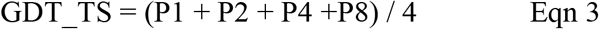

Here, P1, P2, P4, and P8 are the percent of residues with backbone C^α^ RMSD’s < 1 Å, < 2Å, < 4 Å, and < 8 Å, respectively, for consensus reliably modeled / well-define residue ranges of the superimposed structure pairs. GDT_TS = 100% would mean that all consensus reliably modeled residues superimpose with backbone C^α^ RMSD < 1 Å; while GDT_TS of 50% corresponds to an average backbone RMSD of about 4 Å. For brevity, GDT_TS scores are referred to throughout this paper as GDT scores, and are reported as real numbers between 0 and 100.

### Molecular Modeling

Molecular visualization and preparation of graphical representations for figures was done using *PyMol* (DeLano, 2002).

## Results

### RPF-DP and ANSURR scores for assessment of prediction and experimental NMR models in CASP14

In previous studies, we have explored the value of RPF-DP scores in assessing models generated in the Critical Assessment of Protein Structure Prediction (CASP) rounds 13 and 14 (Sala et al., 2019; Huang et al., 2021). When an accurate model is used as the reference structure for computing GDT scores, there is generally a strong correlation between the DP score and GTD (Figure 2A and 2B). In this analysis, consensus well-defined and reliably predicted residue ranges were used for calculating GDT scores between CASP14 NMR structures and prediction models (Table 1), as described previously (Huang et al. 2021). Prediction models from some 100+ prediction groups in each CASP edition, each contributing as many as 5 models, span a wide range of structural accuracy. Some poor prediction models can even return negative DP scores (Figure 2) meaning that the agreement between the model and the chemical shift / NOESY peak list data is poorer than what would be expected for a random coil polypeptide conformation (Huang et al., 2005). Interestingly, for CASP14 target T1055, the best prediction models (provided by AlphaFold2) have DP scores *higher than the experimental NMR structures* (Huang et al., 2021). For target T1027, the DP score is correlated with model prediction accuracy; however the best prediction models (again from AlphaFold2) have DP scores lower than the NMR-derived models. This result is attributable to specific dynamic features of this protein, and are analyzed in detail in Huang et al. (2021). Target T1029 presented an especially instructive case. The initial NMR structure provided for CASP model assessment, T1029_original, had a relatively poor DP score (~ 0.25), and when used as a reference model for the GDT analysis resulted in a poor correlation between DP and GDT across 529 prediction models (Figure 2C, linear correlation coefficient r^2^ = 0.05). Recognizing a potential problem, the experimentalists reassessed their NOESY data for this target, and redetermined the NMR structure. The resulting models have much improved DP scores (~ 0.70), though only marginally better than the AlphaFold2 prediction models, and when used as a reference structure for GDT analysis provide a strong correlation between DP and GDT (r^2^ = 0.87) as expected for an accurate reference structure (*cf* Figures 2C and 2D). These results illustrate the value of DP scores in assessing model accuracy using experimental NOESY data.

**Fig. 2.**
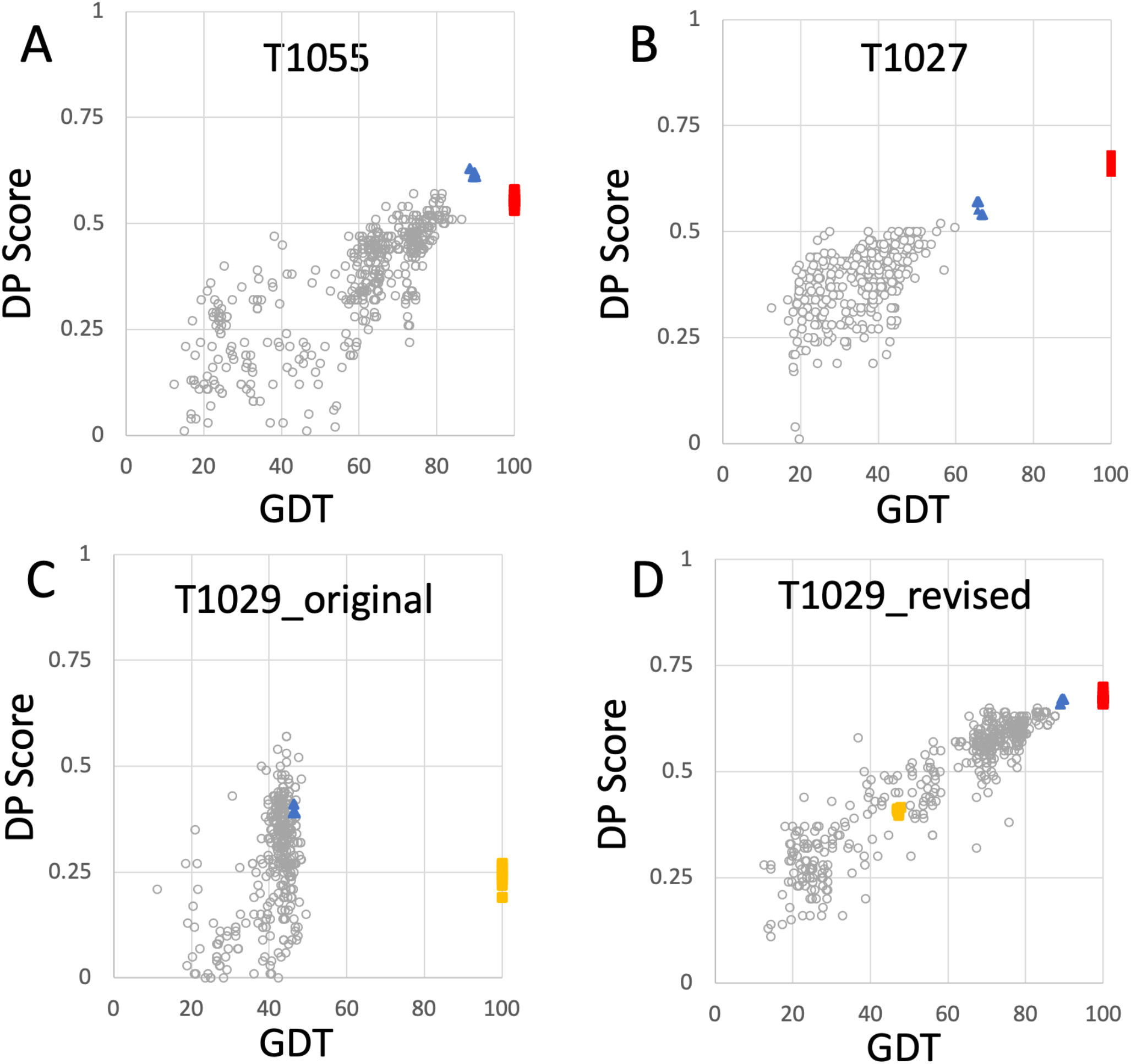
Plots of DP score vs. GDT for NMR and AlphaFold models. For each model, the DP score compares model vs NMR NOESY peak list data, and the GDT score is a measure of similarity to the NMR conformer with best DP score (Huang *et al*., 2021). Plots are provided for (A) target T1055 (511 CASP14 models; linear correlation coefficient r^2^ = 0.66) (B) target T1027 (520 CASP14 models; r^2^ = 0.51) (C) target T1029_original (529 CASP14 models; r^2^= 0.05), and (D) target T1029_revised (529 CASP14 models; r^2^ = 0.87). Open circles are values for CASP14 prediction models (excluding AF models), red squares are the NMR structure models deposited in the PDB, and blue triangles are AF prediction models. In panels C and D, the original NMR structures of target T1029, before revised analysis of NOESY data, are indicated by yellow squares. Negative DP scores are returned for a few models which fit the NMR data more poorly than expected for a random coil conformation (models with DP > −1, not shown). These data are replotted from reference (Huang et al., 2021).

**Table 1.** NMR and X-ray crystal structures used in this study.

| Sample | PDB ID /<br>BMRB ID | PDB<br>Release<br>Date | Total<br>number<br>of<br>residues<sup>a</sup> | Consensus well-defined residue range<sup>b</sup> | Alignment<br>media for RDC<br>measurements |
| --- | --- | --- | --- | --- | --- |
| **CtR107** |  |  |  |  |  |
| X-ray | 3E0H | 2008-09-30 | 158 | 4-158 |  |
| NMR | 2KCU / 16097 | 2009-01-20 |  |  |  |
| **GmR137** |  |  |  |  |  |
| X-ray | 3CWI | 2008-05-06 | 70 | 1-63 |  |
| NMR | 2K5P / 15844 | 2008-08-26 |  |  |  |
| **RpR324** |  |  |  |  |  |
| X-ray | 3LMO | 2010-02-16 | 93 | 4-91 |  |
| NMR | 7TZD / 18263 | this work |  |  |  |
| NMR* | 2LPK / 18263 | 2012-02-29 |  |  | PEG / Phage |
| **SgR42** |  |  |  |  |  |
| X-ray | 3C4S | 2008-02-12 | 58 | 1-56 |  |
| NMR | 2JZ2 / 15604 | 2008-01-22 |  |  |  |
| **SgR209C** |  |  |  |  |  |
| X-ray | 3OSJ | 2010-10-06 | 147 | 13-38, 47-134, 138-143 |  |
| NMR | 2L06 / 17031 | 2010-08-25 |  |  |  |
| NMR* | 7TZ8 / 17031 | this work |  |  | PEG / PAG |
| **SrR115C** |  |  |  |  |  |
| X-ray | 3MA5 | 2010-04-07 |  | 2-92 |  |
| NMR | 2KCL / 16084 | 2009-01-06 | 92 |  |  |
| NMR* | 2KCV / 16821 | 2009-01-20 |  |  | PEG / Phage |
| **<u>CASP14 NMR</u>** |  |  |  |  |  |
| **<u>Targets</u>** |  |  |  |  |  |
| **T1055** | 6ZYC / 34545 | 2021-05-19 | 148 | 310 – 426 |  |
| **T1027** | 7D2O / 36385 | 2020-12-02 | 174 | 36 – 75; 96 – 145 |  |
| **T1029_orig<sup>d</sup>** | 6UF2 / 30676 | 2020-09-30 | 125 | 3-19, 29-46; 53-122 | PAG |
| **T1029_revised<sup>d</sup>** | 7N82 / 30925 | 2021-07-14 | 125 | 3-19, 29-46; 53-122 | PAG |

<sup>a</sup> Number of residues in deposited NMR structure, excluding disordered purification tags, if any.
<sup>b</sup> Consensus of well-defined residues in NMR structures determined by Cyrange (Kirchner and Güntert, 2011) and FindCore2 (Snyder and Montelione, 2005; Snyder et al., 2014), and “reliably modeled” residues determined by AF.
<sup>c</sup> The asterisk (\*) designates the structure was modeled with <sup>15</sup>N-<sup>1</sup>H RDC data using two alignment media.
<sup>d</sup> For target T1029 three kinds of RDC data in one alignment medium were available: N-H<sup>N</sup>, C<sup>α</sup>-C', and C<sup>α</sup>-H<sup>α</sup>

The same CASP14 NMR structures were also assessed using ANSURR (Figures 3 and 4). A similar analysis of CASP14 prediction models has also recently been reported by Fowler and Williamson, the developers of ANSURR (Fowler and Williamson, 2022). Plots of ANSURR corr vs RMSD score (Figure 3) demonstrate the power of ANSURR in identifying accurate prediction models. For CASP14 target T1055, ANSURR scores for the AlphaFold2 models (blue triangles) are somewhat better than for the experimental NMR structures (red squares), consistent with the conclusion of DP analysis. However, many other CASP14 prediction models (with lower GDT to the reference NMR structure) also have very good ANSURR scores (Figure 3A). A similar conclusion can be made for target T1029 (Figure 3C), where the ANSURR scores clearly distinguish the original NMR structure (yellow squares) from the revised NMR structure (red squares). In this case the revised NMR models have somewhat better ANSURR scores than the AlphaFold2 models (blue triangles), also consistent with DP analysis. For target T1027 the ANSURR scores are generally higher for the NMR structures (red squares) than for the AlphaFold2 models (blue triangles; Figure 3B), which is also consistent with the DP analysis of Fig. 2B. However, this target has significant amounts of not-well-defined (Table 1) (apparently flexible) backbone structure (Wu et al., 2020) which can potentially dominate the ANSURR score. This sensitivity of the ANSURR score to removal of not-well-defined regions is evident by comparing Figures 3B (full-length) and 3D (trimmed).

**Fig. 3.**
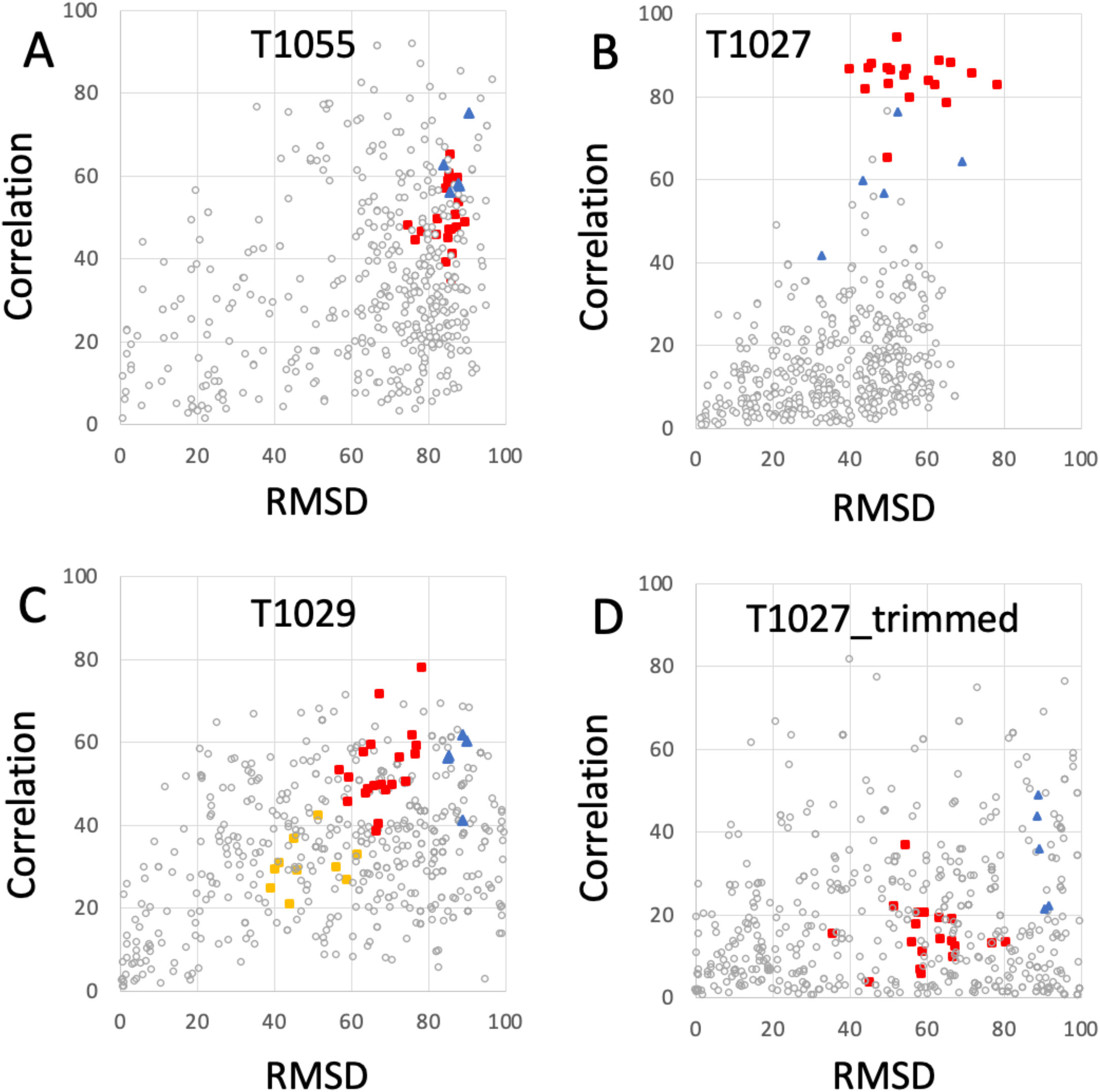
ANSURR Correlation vs RMSD scores for NMR and AlphaFold prediction models. (A) CASP14 target T1055, (B) T1027, (C) T1029 (data shown for both T1029_original and T1029_revised NMR structures), and (D) target T1027_trimmed (residues 36-75 and 96-145) in which coordinates are trimmed to remove the structurally not-well defined (i.e. unreliable) polypeptide segments. As in Fig, 2, in each panel, the open circles are CASP 14 prediction models (excluding AF models), red squares are the final NMR structure models, including T1029_revised, blue triangles are AF prediction models, and yellow squares (in panel C) are for the original NMR structure of target T1029, i.e. T1029_original.

**Fig. 4.**
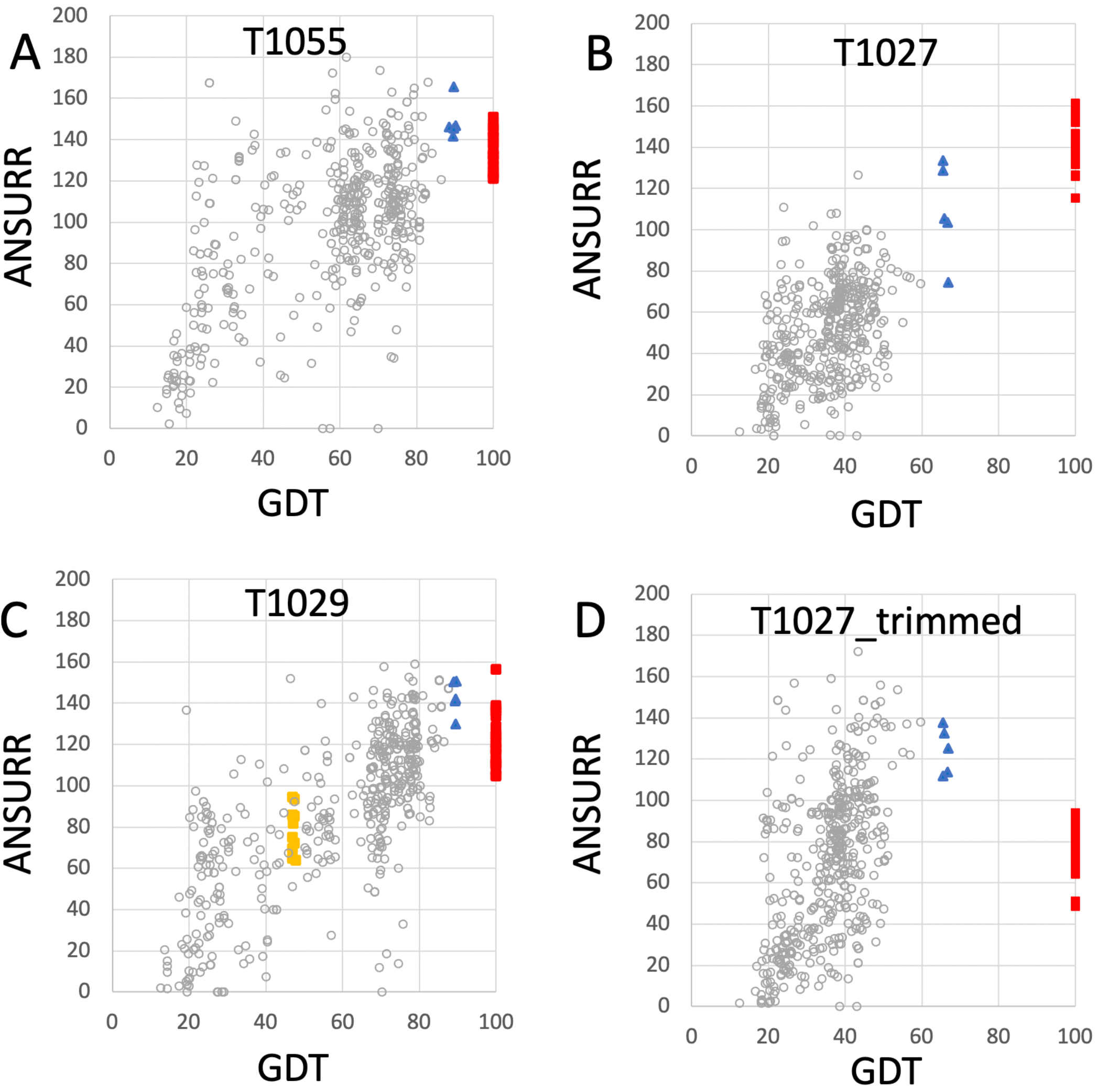
Plots of ANSURR composite score vs. GDT for NMR and AlphaFold models. The data of Fig. 3 were replot to compare the sum of ANSURR correlation and RMSD scores vs. GDT (Huang et al., 2021). Plots are provided for CASP14 targets (A) T1055 (linear correlation coefficient r^2^ = 0.35), (B) T1027 (r2 = 0.47), (C) T1029 (r^2^ = 0.57; data shown for both T1029_original and T1029_revised NMR structures), and (D) T1027_trimmed (residues 36-75 and 96-145, r^2^ = 0.11) in which coordinates are trimmed to remove the structurally not-well defined or unreliably predicted polypeptide segments. In each panel, the open circles are the CASP 14 prediction models (excluding AF models), red squares are the final NMR structure models deposited in the PDB, blue triangles are AF prediction models, and yellow squares are for the original NMR structures of target T1029, before revised analysis of NOESY data.

An important feature of the DP analysis is the correlation between DP and GDT scores when an accurate model is used as the reference structure for calculating the GDT score (Figure 2). Using these CASP14 prediction models, we also assessed if ANSURR scores provide a similar correlation (Figure 4). Generally speaking, the ANSURR scores (corr plus RMSD) do not exhibit as strong correlation with GDT scores as DP scores for these CASP14 NMR targets. The linear correlation coefficients r^2^ for ANSURR (or DP) vs GDT are 0.35 (0.66), 0.47 (0.51), and 0.57 (0.87) for CASP14 NMR targets T1055, T1027, and T1029_revised, respectively. Indeed, some incorrect prediction models with GDT scores as low as 50 have ANSURR scores similar to those of the best AlphaFold2 and NMR models. For the “trimmed” T1027 models, many inaccurate CASP14 prediction models have better ANSURR scores than either the AlphaFold2 or NMR models (Figure 4D). Hence, while ANSURR is a powerful and convenient tool for model quality assessment, requiring only backbone chemical shift data, it is important to complement ANSURR scores with other metrics of structural accuracy.

### NMR /X-ray pairs used for assessment of AF

Six targets from the *NESG NMR X-ray Pairs* data set (Everett et al., 2016) were selected for which NMR structures were determined using standard methods of the NESG Consortium (Liu et al., 2005; Montelione and Szyperski, 2010), but without any residual dipolar coupling data. The PDB id’s (and also BMRB id’s for NMR structures) for these X-ray crystal and NMR structures, together with the PDB coordinate release dates, are listed in Table 1.

The solution NMR structure of target RpR324 is reported in multiple PDB entries, but all of the apo structures were refined with some RDC data. As part of the aim of this study was to assess the accuracy of AF models compared to NMR structures solved with and without RDC data, the deposited chemical shift and NOESY NMR data for RpR324 (BMRB id 18263) were used to re-determine its structure without any RDC data, using our standard NMR structure analysis methods. This structure was deposited in December 2021 as PDB entry 7TZD.

Three of these NESG target proteins were also determined using standard methods that also included refinement with ^15^N-^1^H residual dipolar coupling data (Table 1). For two protein targets, these data were obtained from the *NESG NMR /X-ray Pairs* web site (Everett et al., 2016): RpR324 (2LPK) and SrG115C (2KCV). In a third case, for target SgR209C, ^15^N-^1^H RDC data were available in our database but were not used in the original PDB deposition. For the purposes of this study, the structure was re-determined using the original NOESY, dihedral, and hydrogen bond restraint data (BMRB id 17031) together with these RDC data. This structure and RDC data were deposited in December 2021 as PDB entry 7TZ8.

This process provided 9 solution NMR structures (NMR-based models consisting of ensembles of conformers) for six NESG targets, where all six were determined using chemical shift and NOESY data without ^15^N-^1^H RDC data, and 3 were determined using, in addition, ^15^N-^1^H RDC data. In all three cases where RDC data is available, the RDCs were measured using two alignment media, either polyethylene glycol (PEG) and PF1 phage, or PEG and polyacrylamide stretched gels (PAG) (Table 1).

### Well-defined and not-well-defined regions of models

For structure quality assessment and structural comparisons, it is important to identify “not-well-defined” regions of the protein NMR structure model; i.e. regions where the multiple models (typically 20) generated in the NMR modeling process do not converge, and hence the coordinates are not reliable. Such regions of the structure model will often also have poor knowledge-based structure quality scores and hence are generally excluded from Procheck, Molprobity packing, and Molprobity Ramachandran outlier analysis. Most importantly, not-well-defined regions of NMR structures should not be used for comparisons with prediction models. For identifying not-well-defined regions of these NMR model ensembles, we compared results of Cyrange (Kirchner and Güntert, 2011) and FindCore2 (Snyder and Montelione, 2005; Snyder et al., 2014) (summarized in Supplementary Table S1). These results were generally consistent, and were used to identify consensus ranges of well-defined residue segments. In comparisons with AlphaFold models, we considered only polypeptide segments which are both well-defined in the solution NMR structures and also reliably predicted in the output of AlphaFold; i.e. pLDDT reliability scores > 80 (Jumper et al., 2021b; Jumper and Hassabis, 2022). For the 9 proteins studied here, “not-well-defined” and “not-reliably-modeled” metrics are very consistent with each other (Huang et al, 2021; Supplementary Table S1). The resulting consensus residue ranges of reliably comparable regions are summarized for each protein target in Table 1. The union of the consensus (i) unreliably-modeled residue ranges based on AF pLDDT score and (ii) not-well-defined residue ranges based on NMR structure convergence, were then excluded from the NMR, X-ray, and AF coordinates for structure quality assessment and structure comparisons (e.g. GDT scores). This process “trims” N-terminal and C-terminal regions, as well as some internal polypeptide loop segments. For example, for target SgR209C, this process identified both N- and C-terminal regions, along with two internal polypeptide loops (residue ranges 1 – 12, 39 – 46, 135 - 137, and 144-147, Table 1) for which atomic coordinates are not consistently modeled in the NMR and/or AF models, which were exclude from structural comparisons and knowledge-based structure quality assessment statistics. A similar approach was used to define comparable regions of experimental NMR structures and CASP14 AF prediction models (Huang et al., 2021).

### Modeling with AlphaFold

The six protein targets were modeled with AlphaFold-multimer (Evans et al., 2021) implemented on GPU clusters at RPI, and analyzed with PSVS ver 2.0 (and PDBStat). The resulting well-defined regions of the polypeptide backbone structures are compared for the NMR, AF, and X-ray crystal structures of the six NESG protein targets in Figure 5. In these comparisons, the RDC-refined NMR structure models are shown where available. Generally, the AF, NMR, and X-ray models of these six protein targets are an excellent match. A backbone structural similarity matrix is provided below each superimposed set of backbone structure models (Figure 5). These pairwise comparison scores are between the medoid conformer of each structure ensemble (or the single conformer reported for the X-ray crystal structure). The upper diagonal elements in each matrix are Cα GDT scores, the lower diagonal Cα RMSD scores, and the diagonal values are mean pairwise backbone Cα RMSDs between each member of the structure ensemble and the medoid conformer. In most cases, excluding target CtR107, the NMR, X-ray, and AlphaFold models have high pairwise similarity scores (GDT of 81.5 to 99.6) and low backbone RMSDs (0.48 to 1.68 Å). In particular, the AlphaFold models are in excellent agreement with RDC-refined NMR structures (GDT 91.3 – 99.4). For target RpR324, the AlphaFold models are a better match to the NMR structure, while in the other cases they are a better match to the X-ray crystal structure. For RpR324, the AlphaFold models have high similarity with the NMR structures determined both without (GDT = 92.9, RMSD 1.13 Å) and with (GDT = 99.4, RMSD 0.59 Å) RDCs; the match to the X-ray crystal structure is significantly poorer (GDT = 86.7, RMSD 1.74 Å). The basis of this difference is discussed in more detail below. On the other hand, for target target CtR107 the AlphaFold model is a relatively poor match to the NMR model (GDT = 64.7; RMSD 3.89 Å). The NMR model of CtR107 is particularly poorly converged, with backbone RMSD of well-defined regions within the NMR ensemble of 2.11 Å (compared to the five other targets with more typical ensemble backbone RMSDs of 0.4 to 1 Å). The AlphaFold model of CtR107 is, however, an excellent match to the corresponding X-ray crystal structure (GDT = 97.3; RMSD 1.09 Å).

**Fig. 5.**
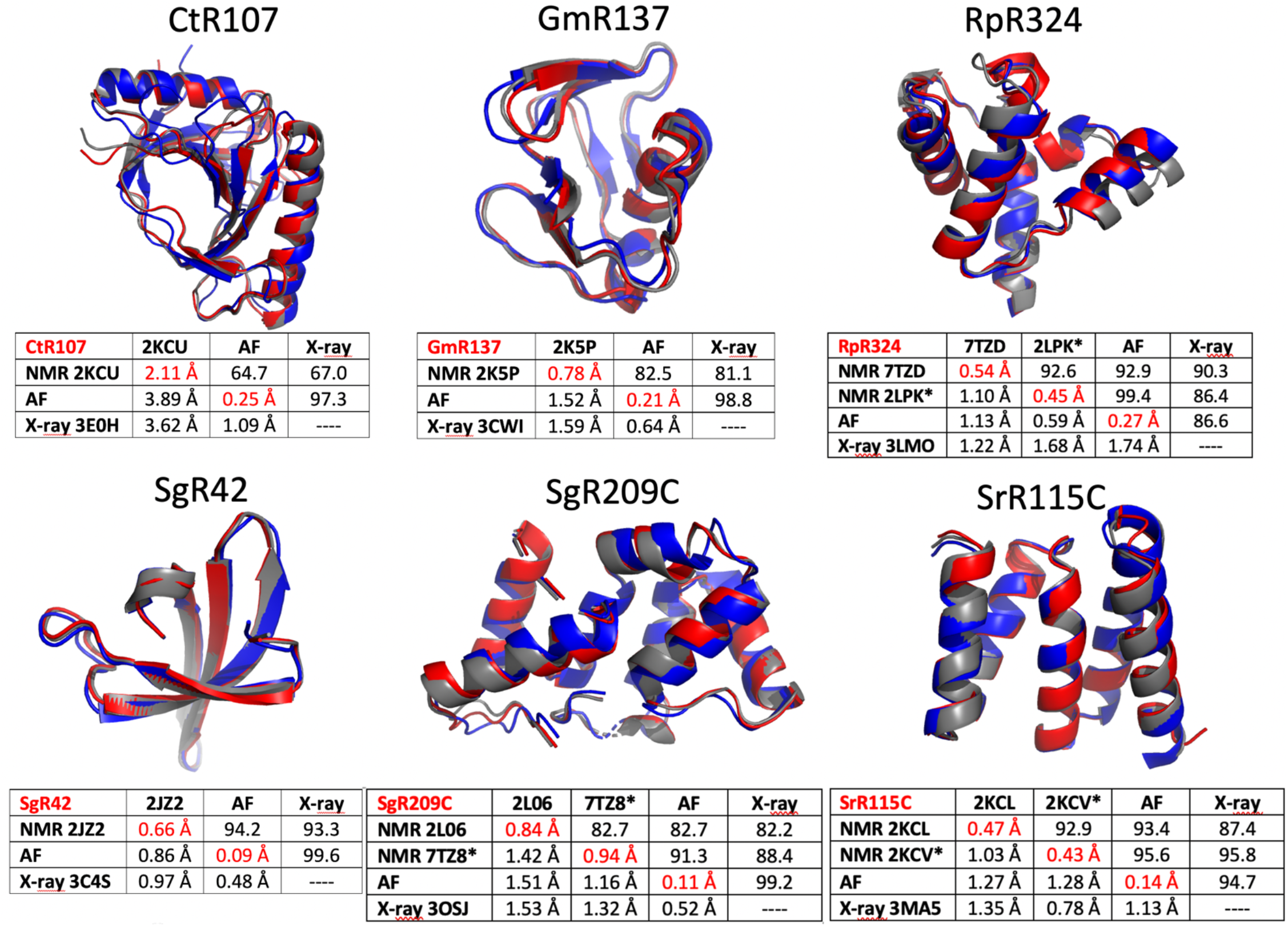
Comparison of AlphaFold, NMR and X-ray crystallography models. Superimposed backbone structures of solution NMR structures (NMR, blue), X-ray crystal structures (X-ray, grey), and AlphaFold prediction models (AF, red) for six proteins selected from the NESG NMR / X-ray pairs database. Below each superimposition is a matrix of backbone structurally-similarity statistics. The upper diagonal provides GDT-TS scores, and the lower diagonal Cα backbone RMSDs. The diagonals (with values in red) are Cα RMSD’s within the corresponding superimposed conformer ensemble relative to the medoid conformer. These models are compared only for residues that are both “well-defined” in the NMR ensemble and “reliably predicted” in the AlphaFold models, as indicated in Table 1. For NMR and AF model ensembles, the coordinates of the medoid conformers are compared. For NMR structures refined with RDC data (i.e., targets RpR324, SgR209C, and SrR115C) the image provided is for the medoid conformer of the structure determined with these RDC data.

### Assessment of NMR and AF models for representative NESG structure using RPF and DP scores

The PSVS v2.0 server was next used to assess <DP> and DP_avg_ metrics (along with R_avg_, P_avg_ and F_avg_ metrics) for the NMR, AlphaFold, and X-ray models (Table 2). Included in this analysis are the corresponding NMR structures refined using a NMR-data-restrained Rosetta modeling protocol (Mao et al., 2014). Generally, for most models and methods, the <DP> score is > 0.70 and DP_avg_ > 0.60, meeting criteria for good quality models (Huang et al., 2012; Huang et al., 2021). *Remarkably, in most cases the AlphaFold models have DP scores (i.e. how well the model fits the NMR data) similar to, or in some cases better than, the NMR structures generated from these same NOESY peak list data*. For each target, the scores for the best performing method is shown in bold font. Considering DP_avg_ as the most discriminating score, compared to the NMR-based models the AF models had the best score (or tied for best score) for three of the six targets; for the other three targets the DP_avg_ score for the AF models was only slightly lower (0.1 units) than the best-scoring NMR-based models. For four of the six targets, the AF models fit the NMR data better than the corresponding X-ray crystal structures; for the remaining two targets the DP_avg_ for the AF models is only slightly lower (0.1 units) than for the corresponding X-ray crystal structure. *Hence, AlphaFold, using no sample-specific experimental data, provides models with an accuracy, based on the DP score, as good or better than the experimentally derived NMR or X-ray crystal structures*.

**Table 2.**
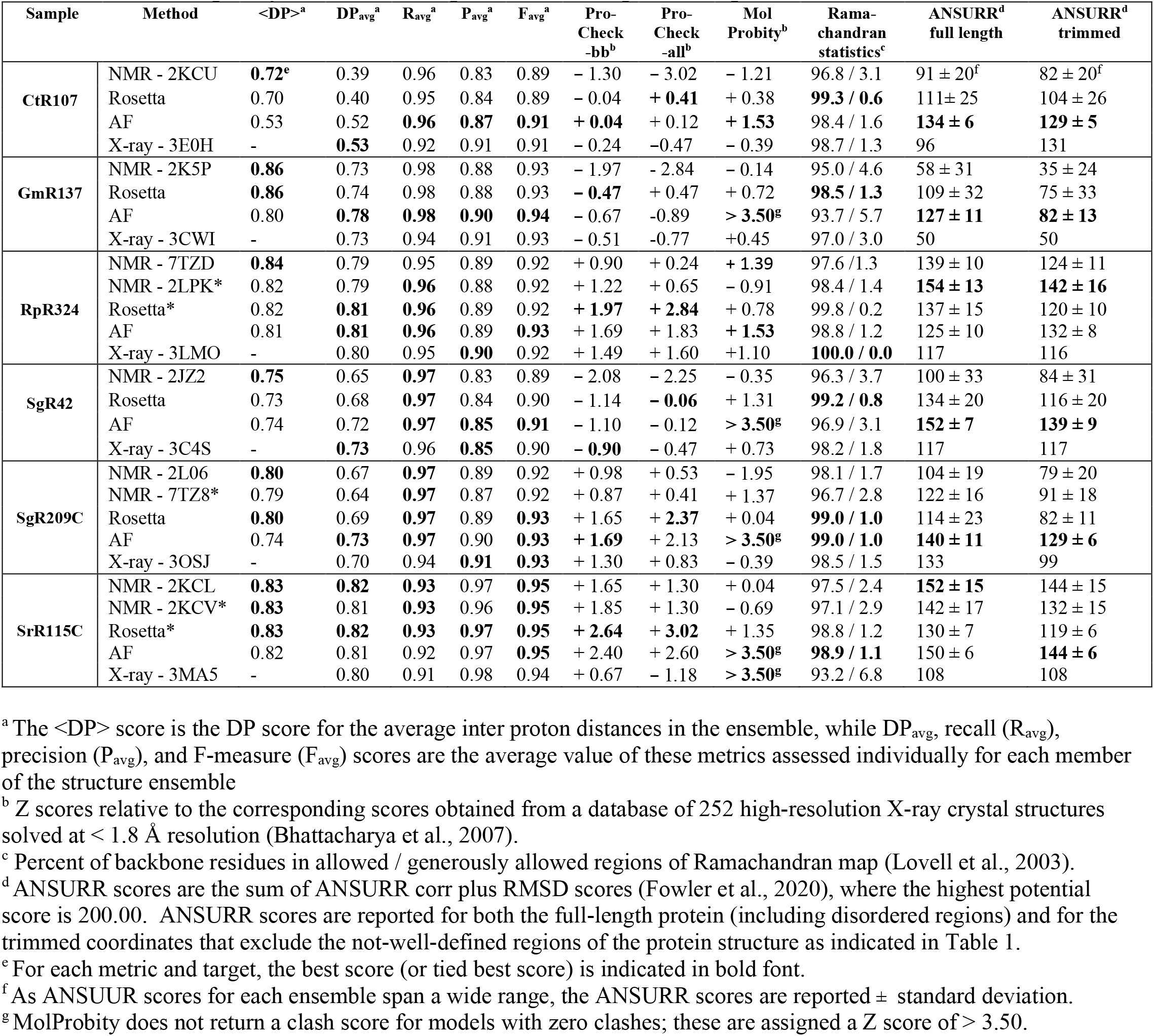
Structure quality statistics for experimental and predicted protein structures.

The one outlier in this analysis is, again, target CtR107. Comparing the <DP> and DP_avg_ provides information about how well individual conformers in the NMR ensemble fit the NMR data. The observation that DP_avg_ is significantly less than <DP> indicates that while the average distances across the (relatively broad) conformer distribution are consistent with the NOESY data, no individual conformer is a good fit to these data. Rather, there appear to be multiple conformations in solution, providing inconsistent NOESY peak data for which no single model is well fit. None of the modeling methods (NMR, AF, or X-ray) provide models with DP_avg_ score > ~ 0.53. This suggests that more powerful ensemble-averaged modeling methods, fitting the data to multiple conformational states, are needed to optimally explain the observed NOESY data obtained for CtR107 in aqueous solution.

### Assessment of NMR and AF models for representative NESG structure using knowledge-based metrics

A necessary, but not sufficient, condition for model accuracy is good knowledge-based structure quality scores. PSVS 2.0 integrates multiple software packages to assess backbone and sidechain dihedral angle distributions using Procheck (Laskowski et al., 1993), and overall packing scores and Ramachandran statistics using MolProbity (Lovell et al., 2003; Chen et al., 2010). ProCheck - backbone and ProCheck - all (backbone plus sidechain) dihedral angle distributions and MolProbity packing scores are normalized to Z scores, such that values of Z > 0 are better quality scores than the mean (Z = 0) obtained for 252 high-resolution X-ray crystal structures (Bhattacharya et al., 2007). These scores assess the quality of buried sidechain conformations in NMR, X-ray, and AlphaFold models. Typically, good models have average Z scores > −3 for these various structure quality scores. For all of the targets and all of the methods, almost all of the models have average Z scores > −3; for many of the targets and methods values of Z are > 0 (Table 2). Ramachandran analysis with Molprobity indicates nearly all backbone dihedral angles are in the allowed and generously allowed phi-psi regions. Generally speaking, the AlphaFold models (and Rosetta-refined NMR models) have excellent knowledge-based dihedral Z score, packing Z scores, and Ramachandran statistics for these metrics; in some cases the AlphaFold models have no Molprobity packing violations at all (in these cases MolProbity does not provide a proper packing score, and the Z score is defined as > +3.5 in Table 2). Both the backbone and core sidechain conformations of the AlphaFold models have excellent knowledge-based validation statistics.

### Assessment of NMR and AF models for representative NESG structure using ANSURR

The models provided for these targets and methods were also assessed by ANSURR (Table 2). Using both full-length and trimmed (as defined in Table 1) models, the best (or second best) ANSURR scores (corr + RMSD) were returned for the AlphaFold models. The one exception was for target RpR324, where the RDC-refined experimental NMR models returned the highest ANSURR score. It is interesting to observe that ANSURR scores have a wide diversity across the models generated by the different modeling methods used for the same protein target, suggesting that it provides strong structural discrimination.

### Assessment of NMR and AF models for representative NESG structure using RDC data

For three of these targets, ^15^N-^1^H RDC data was also available, allowing assessment of models against RDC data. These data are plotted in Figure 6, and the resulting analyses provided by PSVS /PDBStat are summarized in Table 3. For five of the six RDC data sets (three targets, each in two RDC alignment media) the NMR structures determined with the RDC data (with or without Rosetta refinement) were the best fit to the RDC data. For all six RDC data sets, the AlphaFold models fit the RDC data better than the NMR-based models generated from NOE data without RDC data. In five of six cases, the AlphaFold models do not fit the RDC data as well as models determined using these RDC data; however in the sixth case, target SgR209C in PEG alignment media, the AlphaFold models have an even better fit to RDC data (lower Q factor) than the corresponding NMR-derived models generated using these RDC data. The same conclusion is demonstrated by linear regression analysis (R^2^) of calculated vs observed RDC values (Figure 6). *Overall, the AlphaFold models fit the experimental RDC data better than NMR structures generated without RDC data, and in some cases have RDC Q factor and linear regression (R^2^) of calculated vs observed RDC values rivaling those obtained for NMR models refined with these RDC data*.

**Fig. 6.**
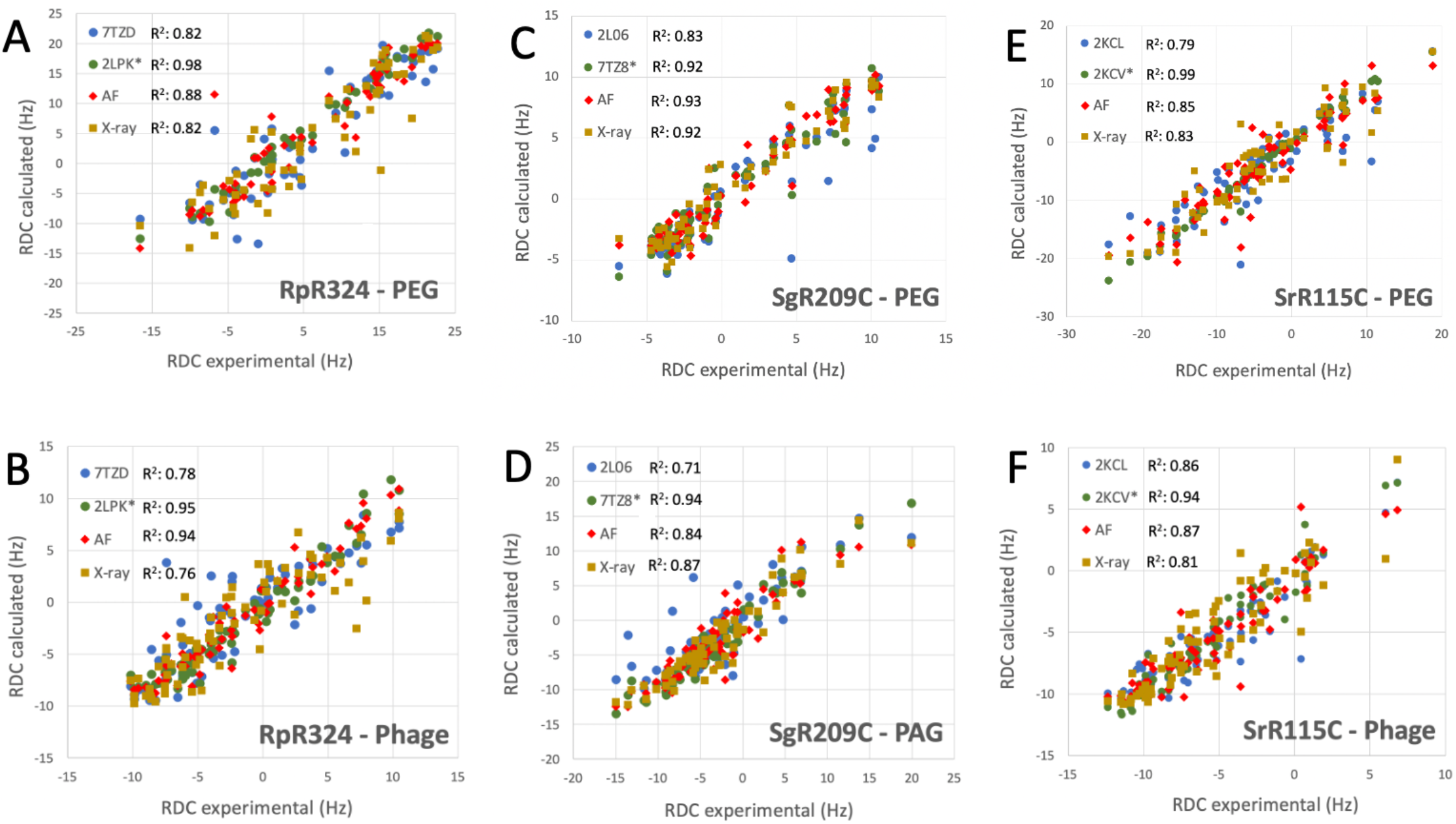
AF structures have excellent fit to RDC data. Comparison of experimentally measured ^15^N-^1^H RDC data (plotted on x-axis) and values computed from experimental or prediction models using PDBStat (Tejero et al., 2013). The data points are for (blue) NMR models determined without RDC data, (green) NMR models refined with ^15^N-^1^H RDC data, (red) AlphaFold prediction models, and (gold) X-ray crystal structures. For NMR and AlphaFold model ensembles, the medoid conformer of the well-defined regions (as indicated in Table 1) were compared. The linear correlation coefficient (R^2^) for each data set is shown in the inset.

**Table 3.** ^15^N-^1^H RDC Q Factors for experimental and AF models^a^.

| Sample | Method | Q1<sup>b</sup> | Q2<sup>c</sup> |
| --- | --- | --- | --- |
| **RpR324**<br>**PEG** | NMR - 7TZD | $0.383 \pm 0.022$ | $0.474 \pm 0.036$ |
|  | NMR - 2LPK<sup>*,d</sup> | **$0.122 \pm 0.011^e$** | **$0.144 \pm 0.014^e$** |
| | Rosetta* | $0.323 \pm 0.016$ | $0.364 \pm 0.023$ |
| | AF | $0.283 \pm 0.021$ | $0.331 \pm 0.032$ |
|  | X-ray - 3LMO | 0.395 | 0.464 |
| **RpR324**<br>**Phage** | NMR - 7TZD | $0.454 \pm 0.042$ | $0.560 \pm 0.074$ |
| | NMR - 2LPK* | $0.225 \pm 0.014$ | **$0.239 \pm 0.016$** |
| | Rosetta* | **$0.220 \pm 0.015$** | $0.245 \pm 0.018$ |
| | AF | $0.227 \pm 0.010$ | $0.253 \pm 0.012$ |
|  | X-ray - 3LMO | 0.459 | 0.564 |
| **SrR115C**<br>**PEG** | NMR - 2KCL | $0.430 \pm 0.024$ | $0.368 \pm 0.033$ |
|  | NMR - 2KCV* | **$0.135 \pm 0.047$** | **$0.111 \pm 0.043$** |
| | Rosetta* | $0.292 \pm 0.019$ | $0.257 \pm 0.025$ |
| | AF | $0.357 \pm 0.007$ | $0.327 \pm 0.010$ |
|  | X-ray - 3MA5 | 0.378 | 0.355 |
| **SrR115C**<br>**Phage** | NMR - 2KCL | $0.250 \pm 0.014$ | $0.319 \pm 0.022$ |
|  | NMR - 2KCV* | **$0.162 \pm 0.016$** | **$0.205 \pm 0.020$** |
| | Rosetta* | $0.170 \pm 0.013$ | $0.218 \pm 0.022$ |
| | AF | $0.235 \pm 0.006$ | $0.321 \pm 0.010$ |
|  | X-ray - 3MA5 | 0.275 | 0.348 |
| **SgR209C**<br>**PEG** | NMR - 2L06 | $0.471 \pm 0.057$ | $0.489 \pm 0.070$ |
| | NMR - 7TZ8* | $0.276 \pm 0.036$ | $0.277 \pm 0.038$ |
| | Rosetta | $0.499 \pm 0.053$ | $0.546 \pm 0.078$ |
|  | AF | **$0.256 \pm 0.009$** | **$0.266 \pm 0.012$** |
|  | X-ray - 3OSJ | 0.272 | 0.307 |
| **SgR209C**<br>**PAG** | NMR - 2L06 | $0.503 \pm 0.065$ | $0.433 \pm 0.073$ |
|  | NMR - 7TZ8* | **$0.243 \pm 0.030$** | **$0.191 \pm 0.028$** |
| | Rosetta | $0.535 \pm 0.061$ | $0.501 \pm 0.094$ |
| | AF | $0.371 \pm 0.010$ | $0.303 \pm 0.012$ |
|  | X-ray - 3OSJ | 0.323 | 0.273 |

<sup>a</sup> $^{15}\text{N}$ - $^1\text{H}$ RDC Q factors were computed for “trimmed” residue ranges shown in Table 1.
<sup>b</sup> RDC Q1 factors computed by the method of Cornilescu et al (Cornilescu et al., 1998).
<sup>c</sup> RDC Q2 factors computed by the method of Clore et al. (Clore and Garrett, 1999).
<sup>d</sup> For each RDC Q factors assessment in each alignment media, the NMR structures refined with the RDC data are indicated with an asterisk (\*). NMR-restrained Rosetta refinement was carried out with $^{15}\text{N}$ - $^1\text{H}$ RDC data for targets SrR115C and RpR324, but did not use RDC data for target SgR209C
<sup>e</sup> The best (lowest) score for each target and method is indicated in bold font.

## Discussion

For the twelve data sets available for nine protein targets, the AlphaFold models have remarkably good fit to the experimental NMR data. Across a wide range of structure validation methods, including both knowledge-based validations of backbone / sidechain dihedral angle distribution and packing scores, and model vs. data validation against experimental NOESY and RDC data, the AF models have similar, and in some cases, better structure quality scores compared with models generated using conventional structure generation methods in the hands of experts using these same NMR data.

The DP score for assessment of NMR derived models has been used routinely in our laboratory, and by various scientists associated with the Northeast Structural Genomics Consortium, as a primary validation tool since its development as a “NMR R factor” in 2005 (Huang et al., 2005; Huang et al., 2006; Raman et al., 2010; Huang et al., 2012; Rosato et al., 2012; Rosato et al., 2013; Rosato et al., 2015; Sala et al., 2019; Huang et al., 2021) (https://montelionelab.chem.rpi.edu/rpf/). However, despite its sensitivity to model inaccuracies and its power for refining NOESY peak list data (Huang et al., 2012), it has not been broadly adopted by the protein NMR community. Here we describe incorporation of RPF-DP analysis into the PSVS 2.0 software package and server. Hopefully this integration will provide broader access to these valuable tools.

In this study we outline the value of using a series of models, generated in this case by the CASP community, to evaluate the correlation between DP and GDT. In this analysis, carried out with good quality NMR data, an accurate reference structure provides a linear correlation (e.g. Fig. 2 A, B, and D), while an inaccurate reference structure provides a poor correlation (Fig 2C). Another valuable insight is provided by comparing the <DP> score, based on the average interproton distance within a model, and DP_avg_, the average DP score computed for each model in the ensemble. In cases where the ensemble is tight and all the models fit the data, these two metrics are similar. However, as observed for the case of target CtR107, for dynamic systems with NOESY data arising from multiple conformations, no single conformer model explains all of the NOESY peak list data to provide a good DP score, and <DP> is much greater than DP_avg_. Such dynamic protein structures also challenge currently available deeplearning-based modeling methods like AlphaFold (Huang et al., 2021).

For most of the cases studied here AlphaFold models returned ANSURR scores similar to or better than the experimentally-determined NMR or X-ray structures (Table 2). ANSURR scores are particularly powerful for protein structure quality assessment since they utilize backbone NMR resonance assignment data that is obtained early in the traditional structure determination process. Chemical shift data are also part of a PDB NMR structure deposition, and is available for many NMR structures. ANSURR scores are sensitive to hydrogen-bonding and accurate atomic packing (Fowler et al., 2020; Fowler and Williamson, 2022), and hence can potentially be improved by energy refinement of the structure model. However, ANSURR score can also be affected significantly by large “not-well-defined” or disordered regions of the protein structure (Fowler and Williamson, 2022), as illustrated in Figures 3 and 4. In addition, for the CASP14 models studied here we observe that there is not a strong correlation between ANSURR and GDT score, potentially complicating the interpretation of the ANSURR score.

Plots of ANSURR vs DP scores for the CASP14 NMR targets (Supplementary Figure S1) have surprisingly low linear correlations (r^2^ ranging from 0.05 to 0.38). This suggests that DP and ANSURR scores provide complementary information for protein NMR structure validation. Indeed, the DP score generally has a better correlation with structural accuracy than the ANSURR score. Since not-well-defined regions often still contribute to the NOESY data, these cannot be excluded from DP analysis. While protein NMR structures deposited in the PDB include chemical shift data, most do not include NOESY peak list (or raw experimental free induction decay NMR data) required for the RPF-DP analysis. Practitioners of protein NMR structure determination using NOE data are encouraged to use DP score analysis for refining atomic models and aid in the accurate interpretation of NOESY peak lists from NMR spectral data.

Our analysis of NMR models generated with and without RDC data revealed that for six data sets for three targets, *AlphaFold models fit these RDC data better than the NMR structures determined without RDC data*. In some cases, the AlphaFold models have RDC Q factors rivaling those obtained for NMR structures determined with these same RDC data. These observations strongly support the concept of using AlphaFold models as a starting point for RDC-based structure determination, without the need for generating NMR NOESY data (Cole et al., 2021).

Analysis of the AF predictions for target RpR324 is of particular interest because the NMR and X-ray crystal structure have notably different structures and also different oligomerization states (Ramelot et al., 2012). A detailed analysis of these structural details is presented in Figure 7. Movement of the α3 helix results in a more ‘open’ hydrophobic cavity in the X-ray structure than in the NMR ‘closed’ conformation (Fig 7A). The NMR sample used for this study was largely monomeric, and size-exclusion chromatography with multiple angle light scattering (SEC-MALS) measurements demonstrate less than 13% dimer in solution (Ramelot et al., 2012). Conversely, in the crystal lattice the protein forms a homodimer with an extensive buried interdomain interface. It remains unclear if there is biological significance for dimerization or for the open conformation, although it seems likely that this represents a functionally relevant conformation. We were interested in finding out whether the AlphaFold prediction would match the NMR or X-ray structure more closely and whether the modeling RpR324 as a homodimer using AlphaFold-multimer would provide a model matching the X-ray crystal structure with a more open conformation.

**Fig. 7.**
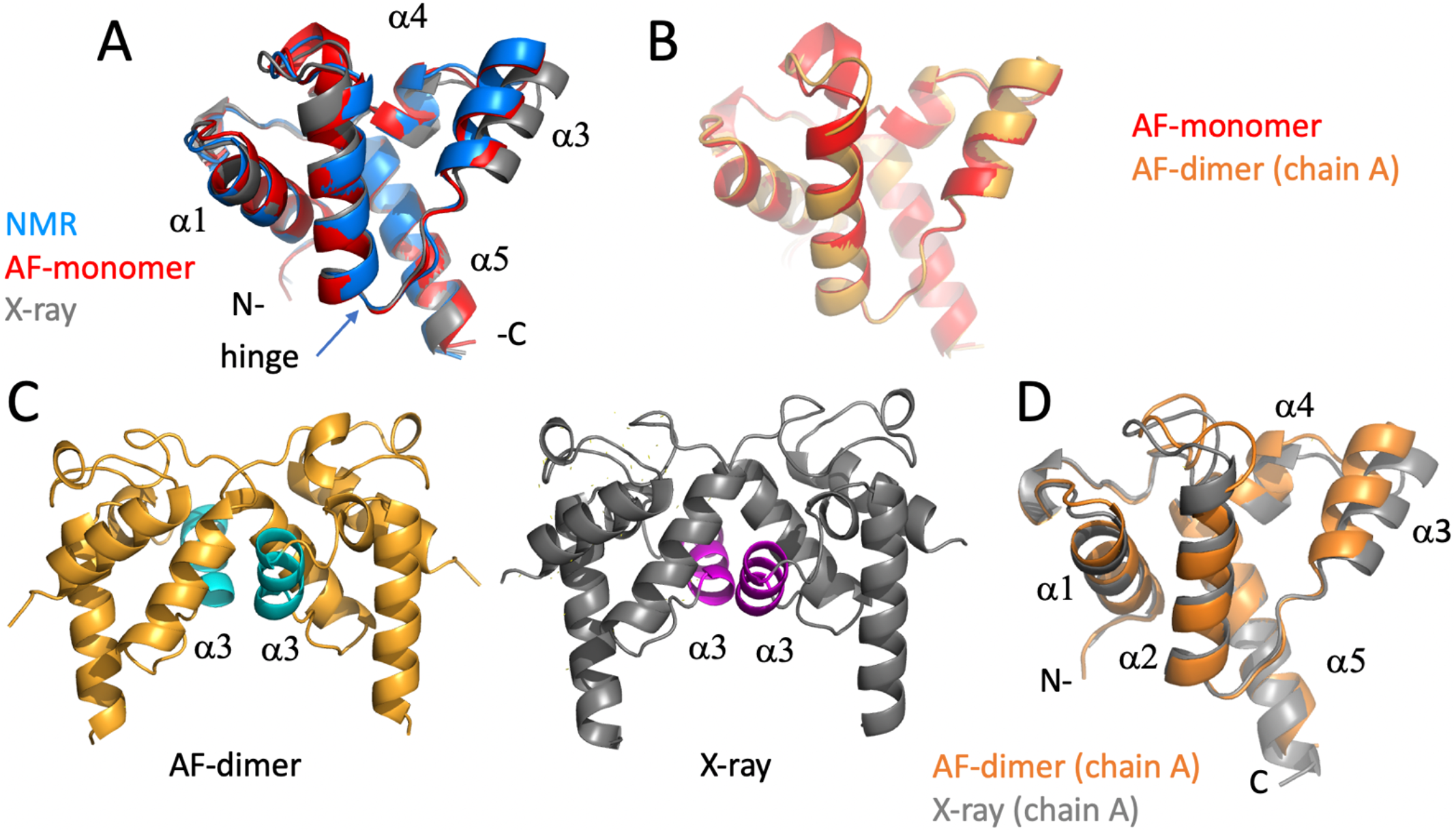
Detailed comparison of solution NMR, X-ray crystal, and AF models of target RpR324. (A) Backbone ribbon representation of solution NMR structure of RpR324 (medoid conformer from PDB ID 2LPK, apo AcpXL) (blue), overlayed with AlphaFold structure calculated as monomer (red), and X-ray crystal structure (PDB ID 3LMO) (grey). (B) Overlay of AlphaFold structure calculated as monomer (red) and calculated as a dimer (orange) using AlphaFold-multimer software. (C) Comparison of (left) AlphaFold dimer (orange) with α3 helix highlighted in cyan (left), with (right) X-ray crystal structure (grey), with α3 helix highlighted in magenta. (D) Overlay of protomers from dimeric AlphaFold model (orange) with X-ray crystal structure (grey), illustrating the significant difference in orientation of α3 helix.

We found that the monomeric AlphaFold models for RpR324 are an excellent match to NMR structures and NMR data, and are significantly better matches to the NMR structure than to the corresponding X-ray crystal structure (Figure 7A). The GDT and RMSD analysis (Figure 5) shows that the AlphaFold model has best structural agreement with NMR structure 2LPK* refined with ^15^N-^1^H RDC data collected with two alignment media (Table 3 and Figure 7A). Based on the RPF-DP analysis, the AlphaFold prediction models agree with the NMR NOESY data just as well as the experimental NMR structure determined using both NOESY and RDC data (2LPK*), and has even better ProCheck and MolProbity clash scores as determined by PSVS analysis (Table 2). The agreement of AlphaFold models with the NMR ^15^N-^1^H RDC data as measured by Q factors is significantly better than that observed for the X-ray structure (Table 3). Taken together, it is clear that the AlphaFold models are more similar to the NMR structures than the X-ray structure, even though the AlphaFold AI was trained on a database of only X-ray structures.

The next noteworthy result is that the dimeric AlphaFold model, generated using AlphaFold multimer, correctly matches the dimer interface observed in the protein crystal, giving further support that this dimerization interface may have biological significance, rather than being a crystallization artifact. The resulting AlphaFold-predicted protomer from this dimer has an almost identical structural match with the monomeric AlphaFold model (Figure 7B), which matches closely the conformation in the monomeric NMR structure (7A). Differences between AlphaFold dimer models and the X-ray crystal structure are still apparent with the ɑ3 helix being more open in the X-ray structure, resulting in small alterations at the dimer interface (Fig 7C, D). Hence, even when modeling the dimeric complex, AlphaFold does not predict the open structure observed in the X-ray crystal structure. The idea that the X-ray structure might represent a functional conformation of RpR324 is, however, supported by the expected biological function of this protein as a specialized acyl carrier protein (AcpXL) for the synthesis of very long-chain fatty acids (20-30 carbons). Although covalent modification of RpR324 on residue S37 by attachment of 4’-phosphopantetheine did not result in any significant changes to the NMR structure, it is believed that addition of a very long chain fatty acid to this carrier arm could favor dimerization or expansion of the hydrophobic cavity as observed in the X-ray structure (Ramelot et al., 2012). While modeling of multiple conformational states of proteins, and energy landscapes, remains an important challenge in the field of protein structure prediction, this structure-function analysis of AcpXL RpR324 illustrates the power of AlphaFold models for developing specific testable hypotheses for driving structural biology research.

One caveat of this study is the fact that the AlphaFold2 AI is trained on X-ray crystal structures available in the PDB through April 2018 (Jumper et al., 2021a). For many of the protein targets predicted in CASP14, including the three CASP14 NMR targets analyzed here, no structures (or structures of homologs) were available in the training data used by AlphaFold2. Although we do not know which X-ray crystal structures were used for training and which were used for testing, the X-ray crystal structures of at least some NESG NMR/X-ray pairs were probably included in the training data. Hence, while these experimental structures (and the structures of homologs) were excluded as templates for AlphaFold modeling, we cannot exclude that there is some kind of indirect memorization within the graph neural network that specifically enhances performance for these targets. However, for about half of the NMR/X-ray pairs we observe DP scores and RDC Q factors indicating that the AlphaFold models fit the NMR data a bit better than the corresponding X-ray crystal structures; this further argues against the notion that the remarkable performance of AlphaFold in generating models that fit NMR data is a trivial result of memorization of specific structural features from the X-ray crystal structures.

Our study begs the question: is experimental NMR structure determination of small, relatively rigid proteins obsolete? Considering the relatively small number of cases studied here, this would be too strong a conclusion. At the very least AlphaFold models need to be validated against experimental data. However, deep learning methods like AlphaFold and RosettaFold have the potential to generate novel insights into structure function relationships at an unprecedented rate, and on genomic and pangenomic scales. Considering this sea change in our ability to generate reliable protein structures, it is important to consider how to use these models to guide sample preparation and data analysis. For example, models could be used for construct optimization, surface analysis for buffer optimization or site-directed mutagenesis to improve spectral quality, interpreting chemical shift perturbations due to protein-ligand, protein-protein, and protein-nucleic acid interactions on modeled protein structures, screening peptides for protein complex formation (Mondal et al., 2022), refining AlphaFold models against RDC, sparse NOE, chemical shift, or paramagnetic NMR data, using models in interpreting NMR data in terms of multiple conformational states of proteins, and the further development of “inverse structure determination” (Huang et al., 2021) in which AlphaFold models are used to guide NMR assignments and data interpretation.

## Conclusions

These studies provide an unambiguous demonstration that AlphaFold can predict structures of small, relatively rigid, single-domain proteins in solution, without structural templates, with an accuracy rivaling experimental NMR studies. AlphaFold models predicted for this study using the platform available in the public domain fit our experimental NMR data (NOEs and RDCs) as well or better than NMR structures generated from these same data by experts using traditional methods. These results contradict the widely held misperception that AlphaFold cannot accurately model solution NMR structures. While AlphaFold and other deep learning methods do not yet have the ability to model multiple alternative conformations of proteins, protein dynamics, and various complex aspects of protein-biomolecular interactions, these models provide a higher starting point with which to begin structure-dynamic-function studies of proteins.

## Abbreviations

AF: AlphaFold
AF2: AlphaFold2
ANSURR: Accuracy of NMR Structures Using RCI and Rigidity
GDT: Global Distance Test
NOE: nuclear Overhauser effect
NOESY: NOE spectroscopy
PAG: PolyAcrylamide Gel (stretched)
PEG: Polythylene Glycol
phage: PF1 phage
pLDDT: predicted Local Difference Distance Test score, a superimpostion independent metric of structure prediction accuracy
PSVS: Protein Structure Validation Software suite
RCI: Random Coil Index for assessing conformational flexibility from chemical shift data
RDC: Residual Dipolar Coupling
RMSD: Root Mean Squared Deviation
RPF-DP score: Recall, Precision, F-measure Discrimination Power score.

## Acknowledgements

RDC data for NESG proteins were recorded in the laboratory of Prof. James Prestegard. We thank all of the scientists of the NESG consortium who produced samples, determined X-ray crystal and NMR structures, and provided these to the community by deposition in the Protein Data Bank. We also thank Prof. M. Williamson, and Drs. N. Fowler, K. Banfa, B. Shurina, and G.V.T. Swapna, for helpful discussions and comments on the manuscript. We also thank Sean Collen for his efforts in porting AlphaFold to the NPL CPU/GPU cluster of the RPI Center for Computing Innovation. This research was supported by grants R01-GM120574 and R35-GM141818 from the National Institutes of Health.

## Author Contributions

RT, YJH, TAR and GTM analyzed data. RT and YJH developed computer codes. TAR determined experimental NMR structures. All authors contributed in writing and editing the manuscript.

