## Supplementary Table S1 and Fig S1 for "AlphaFold Models of Small Proteins Rival the Accuracy of Solution NMR Structures"

**Supplementary Material**

**Supplementary Table S1**. Comparison of well-defined regions in NMR models and reliability-predicted regions of AF models, providing consensus residue ranges for structural comparisons.

| | | Well-define CYRANGE | Well-defined FindCore2 | Reliably Modeled AF pLDDT[a] | Consensus Residue Range |
|---|---|---|---|---|---|
| CtR107 | 2KCU<br>AF | 7-154<br>6-160 | 2-161<br>3-160 | <br>7-62,66-155 | 4-158 |
| GmR137 | 2K5P<br>AF | 1-63<br>1-64 | 1-29,31-64<br>1-67 | <br>1-12,17-63 | 1-63 |
| RpR324 | 7TZD<br>2LPK*,[b]<br>AF | 4-91<br>4-91<br>2-91[c] | 3-9<br>1-93<br>1-98 | 2-92 | 4-91 |
| SgR42 | 2JZ2<br>AF | 2-55<br>1-56 | 1-38,40-56<br>1-60 | <br>1-55 | 1-56 |
| SgR209C | 2L06<br>7TZ8*<br>AF | 14-38,45-143<br>13-39,48-145<br>13-39,46-146 | 13-38,47-143<br>13-39,47-78,83-145<br>12-42,45-146 | 13-41,47-145 | 13-38,47-134,138-143[d] |
| SrR115C | 2KCL<br>2KCV*<br>AF2 | 2-92<br>6-89<br>6-89 | 1-94<br>2-93<br>1-93 | 5-91 | 2-92 |

[a] pLDDT > 80 was used as the cutoff for reliably modeled residue ranges.
[b] NMR models using RDC data are indicated by asterisk (*).
[c] CYRANGE indicates two structural domains: domain 1: 2-55,72-91 and domain 2: 56-71
[d] Rosetta-refined models are not well converged for residue range 135-137.

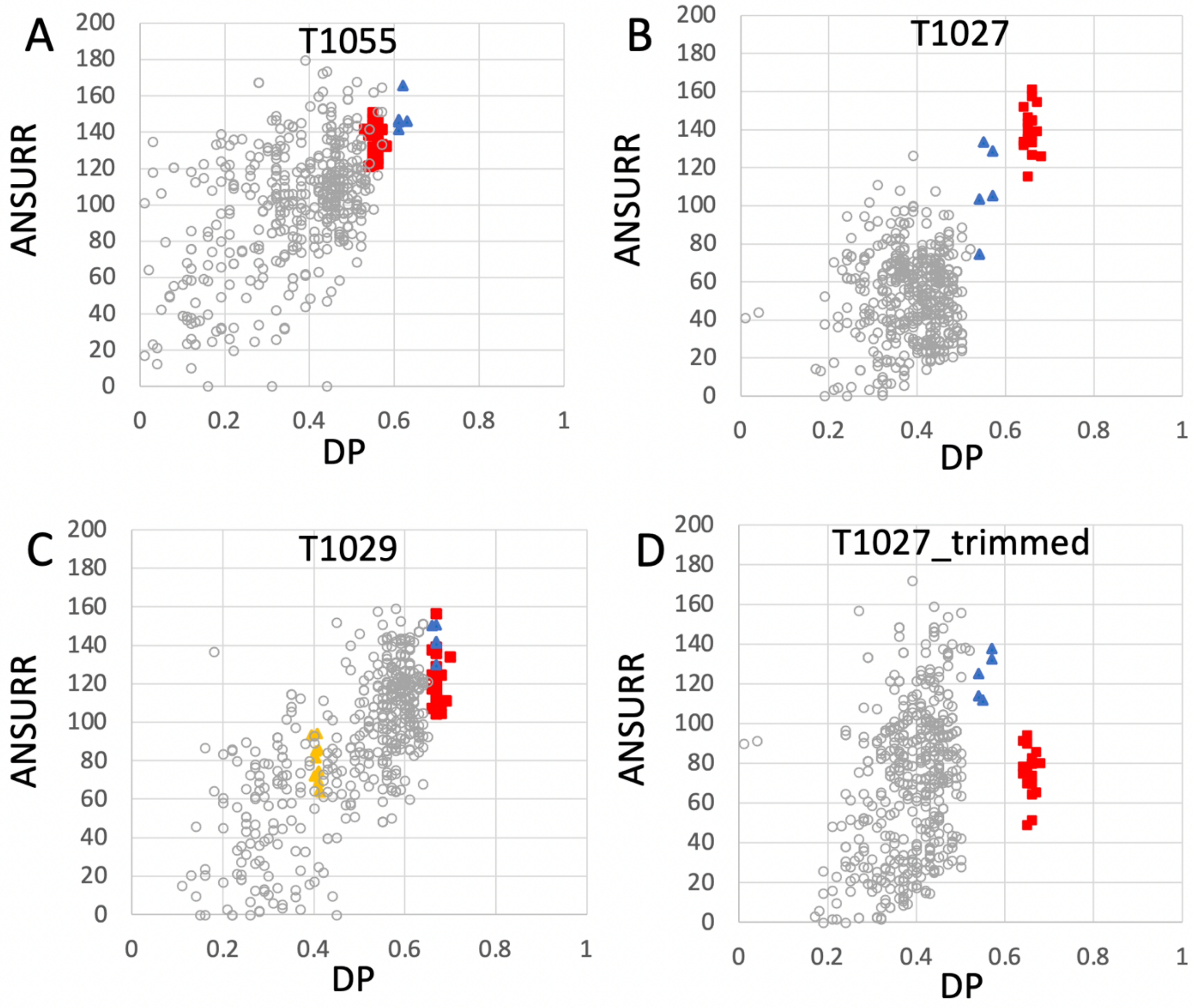


**Supplementary Figure S1. Plots of ANSURR composite score vs. DP score for NMR and AlphaFold models.** Plots are provided for CASP14 targets (A) T1055 (linear correlation coefficient $r^2 = 0.17$), (B) T1027 (r2 = 0.08), (C) T1029 ($r^2 = 0.38$; data shown for both T1029_original and T1029_revised NMR structures), and (D) T1027_trimmed (residues 36-75 and 96-145, $r^2 = 0.05$) in which coordinates are trimmed to remove the structurally not-well defined or unreliably predicted polypeptide segments. In each panel, the open circles are the CASP 14 prediction models (excluding AF models), red squares are the final NMR structure models deposited in the PDB, blue triangles are AF prediction models, and yellow squares are for the original NMR structures of target T1029, before revised analysis of NOESY data.
